# Antxr2-mediated fine-tuning of Collagen VI ensures skeletal muscle function

**DOI:** 10.1101/2025.09.11.675515

**Authors:** Samuele Metti, Lucie Josepha Bracq, Matteo Signor, Nicola Facchinello, Lisa Gambarotto, Leonardo Nogara, Giulia Pigato, Laurence Abrami, Béatrice Kunz, Julien Duc, Patrizia Sabatelli, Aurora De Acutis, Gabriele Maria Fortunato, Giovanni Vozzi, Paola Braghetta, Bert Blaauw, F. Gisou van der Goot, Paolo Bonaldo

**Affiliations:** Department of Molecular Medicine, University of Padova, Padova, Italy; Global Health Institute, School of Life Sciences, Ecole Polytechnique Fédérale de Lausanne, Lausanne, Switzerland; Neuroscience Institute, Italian Research Council (CNR), Padova, Italy; Department of Biomedical Sciences, University of Padova, Padova, Italy; Department of Pharmaceutical and Pharmacological Sciences, University of Padova, Padova, Italy; CNR, Institute of Molecular Genetics “Luigi Luca Cavalli-Sforza”, Unit of Bologna, Bologna, Italy; Department of Information Engineering and Research Centre “E. Piaggio”, University of Pisa, Pisa, Italy

**Keywords:** ECM remodelling, CMG2, muscle hypertrophy, fibro-adipogenic precursors, intracellular ECM degradation

## Abstract

Tissue function relies on the extracellular matrix (ECM) that surrounds cells, providing structural and biochemical support. The complex ECM composition depends on an adequately tuned balance between the deposition and degradation of each of its components. Disequilibrium may cause disease, as observed for Collagen VI (COL6), for which mutations lead to muscular dystrophy. Here, we investigated the role of Anthrax Toxin Receptor 2 (ANTXR2/CMG2), a receptor that controls the turnover of COL6, in skeletal muscle. We show that ANTXR2 is mostly expressed by fibro-adipogenic precursors and that its deficiency in ANTXR2 null (*Antxr2^-/-^*) mice leads to a premature and irregular COL6 accumulation in intramuscular connective tissue. This results in tissue stiffening and gradual, non-functional muscle hypertrophy, marked by impaired locomotion and myopathic signs. Our findings further indicate that COL6 accretion drives these alterations, as revealed by *Antxr2^-/-^::Col6a1^-/-^* double knockout mice, highlighting the essential role of ANTXR2-mediated COL6 remodeling in maintaining ECM homeostasis and muscle functionality.

**SIGNIFICANCE STATEMENT:** Remodelling of the extracellular matrix (ECM) was long thought to rely almost exclusively on extracellular proteases. Increasing evidence, however, indicates that some ECM components may undergo intracellular degradation following receptor-mediated endocytosis, as we have found for Collagen VI (COL6). Here, we identify the COL6 receptor ANTXR2 as a critical regulator of ECM turnover in skeletal muscle.. When ANTXR2 is absent, COL6 builds up, followed by the accumulation of fibrillar collagens, without changes in gene expression. These alterations in muscle ECM lead to increased stiffness, myofiber defects and impaired locomotor activity. Our findings establish ANTXR2 as a key regulator in ECM remodelling, offering new insights into potential treatments for conditions associated with defective ECM remodeling, such as aging and congenital muscular dystrophies.

## INTRODUCTION

The extracellular matrix (ECM), an intricate meshwork of proteins and proteoglycans, is required to organize cells into tissues and maintain organ structure. Such macromolecular network regulates a number of cellular functions and processes through biochemical and biomechanical cues^1,2^. The ECM is highly dynamic, being remodelled according to specific needs through a fine-tuned balance of protein synthesis and assembly, post-translational modifications, recycling and degradation. Failure of this tight regulation contributes to a range of disease conditions such as cancer progression, osteoarthritis, and muscular dystrophies^3–5^. The mechanisms underlying the control of ECM remodelling in the different tissues are however incompletely understood.

ECM degradation can occur through two main routes^5,6^. The first and most documented is extracellular degradation, which is mainly orchestrated by different families of secreted proteases whose activity is responsible for the cleavage of ECM components within the extracellular space^7,8^. The second mode of degradation is intracellular, following receptor-mediated endocytosis of ECM macromolecules and lysosomal clearance^9,10^. Intracellular degradation has been largely overlooked due to the limited characterization of the involved molecular players. In this study, we have investigated the relevance of receptor-mediated clearance of Collagen VI (COL6) in skeletal muscle, given the key role of this ECM component for proper muscle function.

In past work, we identified Anthrax Toxin Receptor 2 (ANTXR2), also known as Capillary Morphogenesis Gene 2 (CMG2), as a cell surface receptor involved in the homeostasis of COL6, by mediating COL6 endocytosis and lysosomal delivery^11,12^. The ANTXR2 name stems from the well-known role of this surface protein in binding and internalizing *Bacillus anthracis* toxins^13^. The physiological roles of ANTXR2 are only beginning to be elucidated^14^. Loss-of-function mutations in the human *ANTXR2* gene cause hyaline fibromatosis syndrome (HFS, MIM #228600)^15^, a severe and debilitating autosomal recessive disease characterized by the formation of non-cellularized subcutaneous and visceral hyalinized nodules highly enriched in COL6^11^. This human condition is mirrored to some extent by ANTXR2 null (*Antxr2^-/-^*) mice, which progressively accumulate COL6 in the uterus, affecting organ contraction, thus leading to the inability to deliver pups^11,16^.

COL6 is made of three major polypeptide chains [α1(VI), α2(VI), α3(VI)] coded by separate genes (*COL6A1*, *COL6A2* and *COL6A3* in humans)^17^. It is unique within the collagen superfamily because it forms a distinctive network of beaded microfilaments in the ECM of various organs, where it binds several ECM components, such as fibrillar collagens^18^, fibronectin^19,20^ and proteoglycans^21^. The proper deposition of COL6 in the ECM is particularly critical for skeletal muscle, and COL6 deficiency due to mutations in its genes is causative for inherited muscle disorders known as COL6-related myopathies^22–24^. COL6 synthesis and matrix deposition are finely regulated in different tissues^25–27^, however the mechanisms that control its turnover in muscle are largely unknown.

In this study, we uncover a critical role of ANTXR2 in regulating COL6 turnover in skeletal muscle by exploiting an ANTXR2-deficient mouse model. We find that ANTXR2 is predominantly expressed by fibro-adipogenic precursors (FAPs) and that its genetic ablation leads to an early onset and progressive buildup of COL6 in muscle ECM, with a subsequent increase in fibrillar collagens accompanied by myofiber changes. Notably, although *Antxr2^-/-^* muscles display hypertrophy, the added muscle mass is non-functional and exhibits hallmarks of myopathy. To determine causality, we generated and performed phenotypic characterization of *Antxr2^-/-^::Col6a1^-/-^* double knockout mice, which provided a genetic proof that excessive COL6 is the key driver of the pathological features observed in *Antxr2^-/-^* skeletal muscle. This work demonstrates that the proper regulation of COL6 homeostasis by ANTXR2 is critical for preserving ECM features and, consequently, muscle structure and function.

## RESULTS

### *Antxr2* is expressed by FAPs in adult skeletal muscle

We first evaluated the expression of *Antxr2* gene in the different cell populations of adult mouse muscles. We mined the GSE147127 dataset containing single-nucleus RNA-seq data of skeletal muscles from C57BL/6 mice of different ages^28^. Analysis of tibialis anterior (TA) muscles from 5-month-old mice revealed that *Antxr2* expression is high in FAPs, whereas it is extremely low in the nuclei of myofibers, also know as myonuclei (Fig. 1A). Expression in FAPs and very low expression in myofibers was also observed for the three major COL6 genes, *Col6a1*, *Col6a2* and *Col6a3* genes (Fig. 1A), consistent with previous findings that they are predominantly expressed by muscle fibroblasts, rather than by mature myofibers^27–29^.

**Figure 1.**
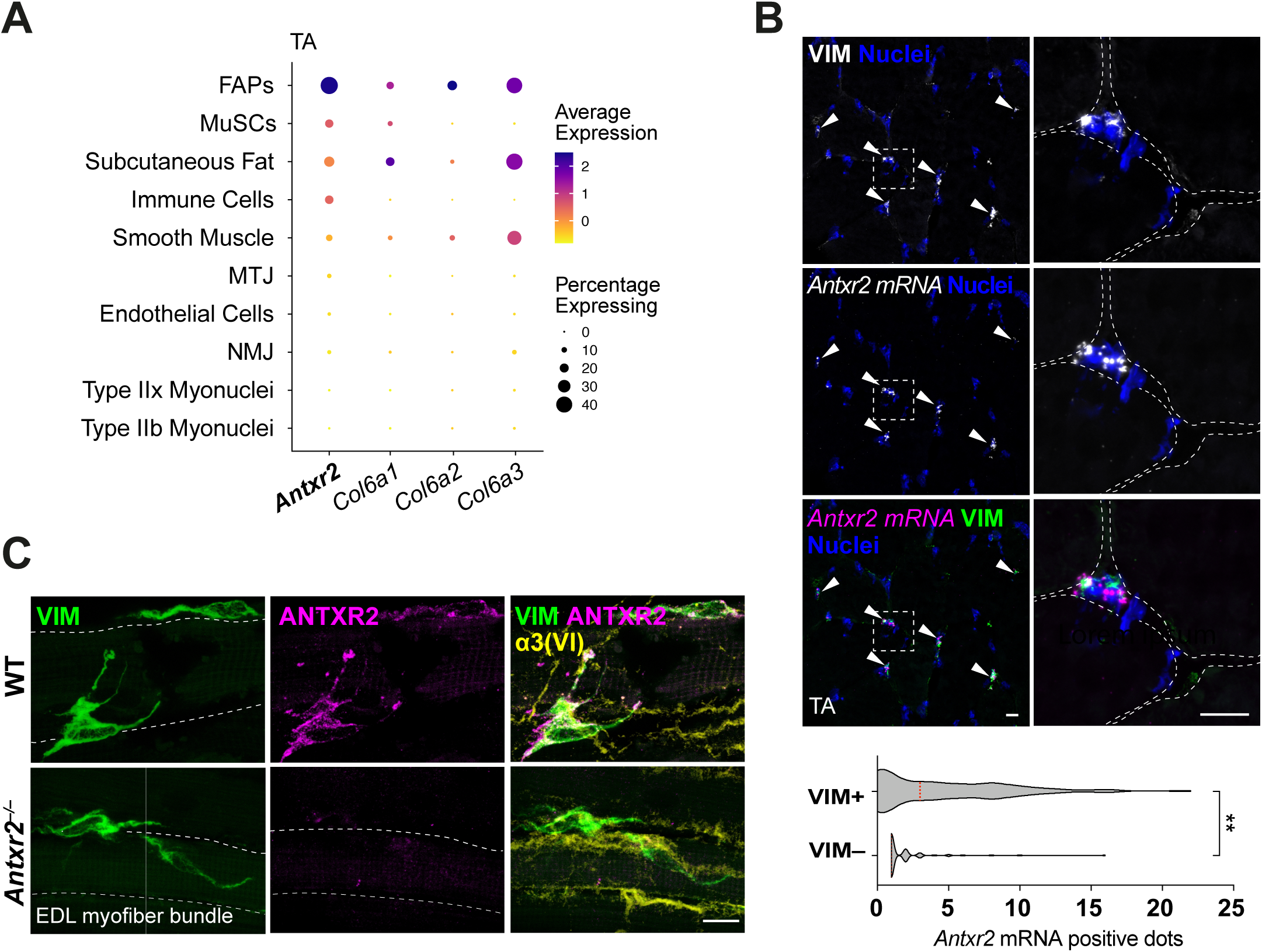
ANTXR2 is mainly expressed by FAPs in adult skeletal muscle. (**A**) Dot plot showing the expression of *Antxr2*, *Col6a1*, *Col6a2* and *Col6a3* genes in the different cell populations of TA muscles from 5-month-old C57BL/6 WT mice, based on available single-nucleus RNA-seq data deposited in the GSE147127 dataset. Dot size reflects the percentage of cells with a non-zero expression level and color scale represents the average expression level across all cells within the cluster. MTJ, myotendineous junction; MuSCs, muscle stem cells; NMJ, neuromuscular junction. (B) Representative fluorescence microscopy images of *in situ* hybridization for *Antxr2* mRNA in TA cross-sections of 8-month-old WT animals, showing colocalization of *Antxr2* mRNA (magenta) with the FAP marker Vimentin (VIM, green). Nuclei were stained with Hoechst (blue). Muscle fibers are outlined with white dashed lines. White arrowheads point at Vimentin-positive cells. The violin plot shows the number of *Antxr2* mRNA positive dots localized in close proximity of Vimentin positive (VIM+) or negative (VIM–) regions (*n* = 20 fields from 3 WT mice; **, *P* <0.01; two-tailed Mann-Whitney test). Scale bar, 10 μm. (**C**) Representative confocal images of immunostaining for Vimentin (VIM, green), ANTXR2 (magenta), and COL6 α3 (α3(VI), yellow) in EDL myofiber bundles of 8-month-old WT and *Antxr2^-/-^*animals. Muscle fibers are outlined with white dashed lines. Scale bar, 10 μm.

To corroborate the RNA-seq data, we performed RNA *in situ* hybridization in TA cross-sections from 8-month-old wild-type (WT) animals. *Antxr2* mRNA was indeed predominantly found in FAPs, positive for the fibroblast marker Vimentin (Fig. 1B). Immunofluorescence of myofiber bundles isolated from Extensor Digitorum Longus (EDL) muscles confirmed ANTXR2 staining in Vimentin-positive FAPs in WT samples, whereas no ANTXR2 signal was detectable in *Antxr2^-/-^* samples (Fig. 1C). Staining with an antibody against the α3(VI) polypeptide showed COL6 localization in the external layer of the basement membrane (Fig. 1C). No signal for ANTXR2 protein was found in WT myofibers (Fig. 1C), nor were any *Antxr2* mRNA-positive dot detected in the proximity of myonuclei (Supplementary Fig. 1). Together, these data show that FAPs are the major cell population expressing ANTXR2 within adult skeletal muscle.

### Abnormal progressive buildup of COL6 in ANTXR2-deficient muscles

Given the ability of ANTXR2 to bind COL6 and mediate its endocytosis and degradation in lysosomes in a primary fibroblast culture system^11,12^, we assessed COL6 abundance in muscles of WT and *Antxr2^-/-^* mice at different postnatal ages. Western blotting for the COL6 α1(VI) and α3(VI) chains in protein lysates of gastrocnemius muscle from 5-day-old pups did not reveal any significant differences between WT and *Antxr2^-/-^* samples (Fig. 2A). To corroborate these findings, we performed immunofluorescence for the COL6 α3(VI) and α6(VI) chains, taking into account previous studies which showed that the α3(VI) polypeptide can be substituted by other minor COL6 chains in some conditions and tissues, and in particular α6(VI) in skeletal muscles^30,31^. For both chains, immunostaining was similar between WT and *Antxr2^-/-^* muscles, with COL6 α3(VI) restricted to perimysium (the thicker connective tissue layer encasing muscle fiber bundles), whereas COL6 α6(VI) was also abundant in the endomysial layer surrounding individual myofibers (Fig. 2B). These biochemical and immunohistochemical data indicate that ANTXR2 ablation does not visibly affect COL6 abundance and localization in skeletal muscle during early postnatal life.

**Figure 2.**
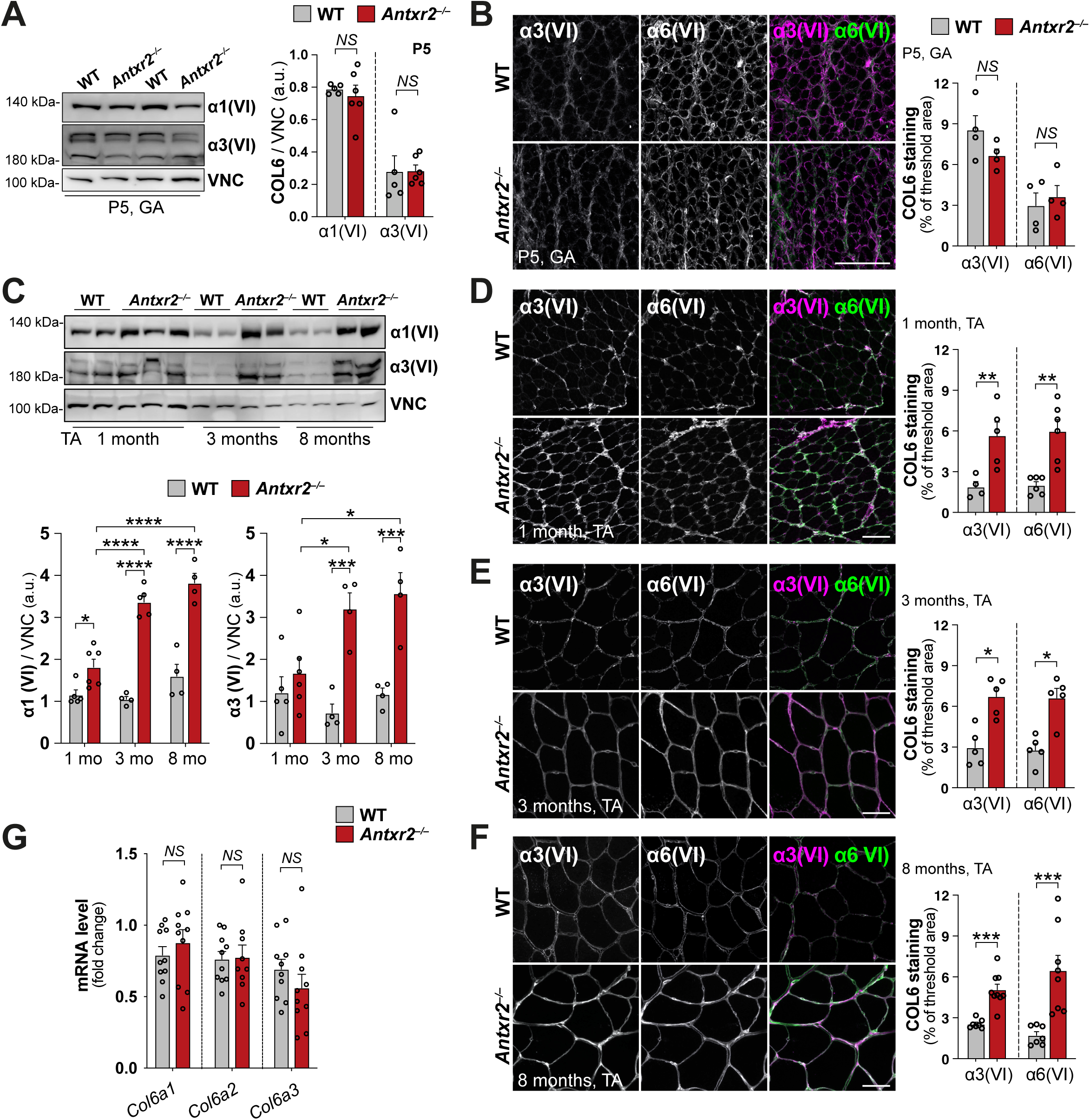
COL6 undergoes progressive accretion in ANTXR2-deficient muscles. (**A**) Western blotting for COL6 α1 [α1(VI)] and α3 [α3(VI)] chains in protein extracts from gastrocnemius muscles of 5-day-old (P5) WT and *Antxr2*^−/−^ mice. Vinculin was used as a loading control. The histograms on the right show the densitometric quantification of α1(VI) and α3 (VI) signals, normalized on Vinculin protein levels. Data are expressed as mean ± s.e.m. (*n* = 5-6, each group; NS, not significant; two-tailed Mann-Whitney test). (**B, D-F**) Representative confocal images of immunostaining for COL6 α3 (α3(VI), magenta) and α6 (α6(VI), green) chains in cross-sections of gastrocnemius muscles from 5-day-old WT and *Antxr2*^−/−^ mice (**B**), and in cross-sections of TA muscle from 1-month-old (**D**), 3-month-old (**E**), and 8-month-old (**F**) WT and *Antxr2*^−/−^ mice. Note that, as expected, myofiber size increases after birth and along early postnatal life, being larger in 3-and 8-month-old when compared to 5-day-old muscle sections. The respective histograms on the right show the percentage of threshold area positive for α3(VI) or α6(VI) staining. Scale bars, 50 μm. Data are expressed as mean ± s.e.m. (*n* = 4–8, each group; NS, not significant; *, *P* <0.05; **, *P* <0.01; ***, *P* <0.001; two-tailed Mann-Whitney test). (C) Western blotting for COL6 α1 [α1(VI)] and α3 [α3(VI)] chains in protein extracts from TA muscles of 1-, 3-and 8-month-old (mo) WT and *Antxr2*^−/−^ mice. Vinculin was used as a loading control. The histograms on the bottom show the densitometric quantification of α1(VI) (left panel) and α3(VI) (right panel) signals, normalized on Vinculin protein levels. Data are expressed as mean±s.e.m. (*n* = 4–6, each group; *, *P* <0.05; ***, *P* <0.001; ****, *P* <0.0001; two-way ANOVA test, with Sidak’s test for multiple comparisons). (**G**) RT-qPCR analysis for *Col6a1*, *Col6a2* and *Col6a3* transcript levels in gastrocnemius muscles of 8-month-old WT and *Antxr2*^−/−^mice. *Gapdh* was used as housekeeping gene. Data are expressed as mean ± s.e.m. (*n* = 9–10, each group; NS, not significant; two-tailed Mann-Whitney test). a.u., arbitrary units; VNC, vinculin; GA, gastrocnemius.

Conversely, at 1 month of age *Antxr2^-/-^* mice displayed a progressive accumulation of COL6 chains, as revealed by western blotting of TA muscles, becoming increasingly more abundant in 3-and 8-month-old mice when compared to the corresponding samples from age-matched WT animals (Fig. 2C). Immunofluorescence of TA muscles at the same postnatal ages confirmed markedly increased of both α3(VI) and α6(VI) in the endomysium of *Antxr2^-/-^* animals (Fig. 2D-F). A similar pattern of COL6 accumulation was displayed by the Gastrocnemius of 8-month-old *Antxr2^-/-^* mice (Supplementary Fig. 2A, B).

To assess whether COL6 protein accumulation was a consequence of increased gene expression, we quantified transcript levels. RT-qPCR analyses in TA of 8-month-old animals showed no significant difference in *Col6a1*, *Col6a2* and *Col6a3* mRNA levels between WT and *Antxr2^-/-^* mice (Fig. 2G). We next considered whether COL6 assembly and secretion could be affected. Past studies demonstrated that COL6 undergoes a distinctive stepwise intracellular assembly process, in which three chains associate into monomers (∼500 kDa), dimers (∼1,000 kDa) and tetramers (∼2,000 kDa), which are finally secreted and deposited in the ECM^17^. Protein electrophoresis into composite acrylamide-agarose gels under non-reducing conditions showed that all COL6 species increased proportionally in TA lysates of *Antxr2^-/-^* mice, indicating that the assembly of monomers, dimers and tetramers is not appreciably affected (Supplementary Fig. 2C). Interestingly, the high COL6 content of *Antxr2^-/-^* muscles led to a significant increase in the SDS-insoluble COL6 fraction (Supplementary Fig. 2D), suggestive of increased crosslinking.

Altogether, these data show that ANTXR2 null muscles undergo a progressive buildup of COL6 in the ECM, despite normal expression of COL6 genes and proper COL6 biosynthesis with the expected formation of monomers, dimers and tetramers. The greatly increased abundance of COL6 and its increased SDS-insolubility in *Antxr2^-/-^* samples therefore support a critical role for ANTXR2 in regulating COL6 turnover in skeletal muscle.

### Fibrillar collagens accumulate in muscles of adult ANTXR2-deficient mice

We analyzed the tissue architecture of TA muscles from 8-month-old *Antxr2^-/-^* mice and found that they did not overtly differ from that of WT mice (Fig. 3A). In particular, histological analysis by haematoxylin-eosin staining did not reveal any sign of fibrous-adipose replacement or cellular infiltration, which would be expected in the presence of active fibrosis or necrotic lesions^32^. However, *Antxr2^-/-^*muscles repeatedly displayed the presence of cytoplasmic inclusions within myofibers, which were not present in the corresponding WT samples (Fig. 3A, white arrowheads). The histological profile of such abnormalities is reminiscent of tubular aggregates, one of the most recurrent myofiber alterations found in neuromuscular diseases^33,34^. This was confirmed by Gomori trichrome staining, which revealed a markedly increased incidence of myofibers containing staining-positive aggregates in 8-month-old *Antxr2^-/-^* male mice when compared to age-and sex-matched WT samples (Fig. 3B, C), predominantly affecting larger glycolytic myofibers. Electron microscopy revealed that these cytoplasmic inclusions were membranous accumulations characterized by tubular and saccular dilations, thereby definitely identifying them as tubular aggregates (Fig. 3D). These stained positively for the sarco/endoplasmic reticulum markers Calsequestrin, STIM1 and SERCA1/2/3, and the T-tubule markers Orai1 and DHPR (Supplementary Fig. 3A-C). Altogether, this analysis shows that TA muscles from 8-month-old *Antxr2^-/-^* mice overal maintain a proper architecture, but display signs of alterations as exemplified by the presence of tubular aggregates.

**Figure 3.**
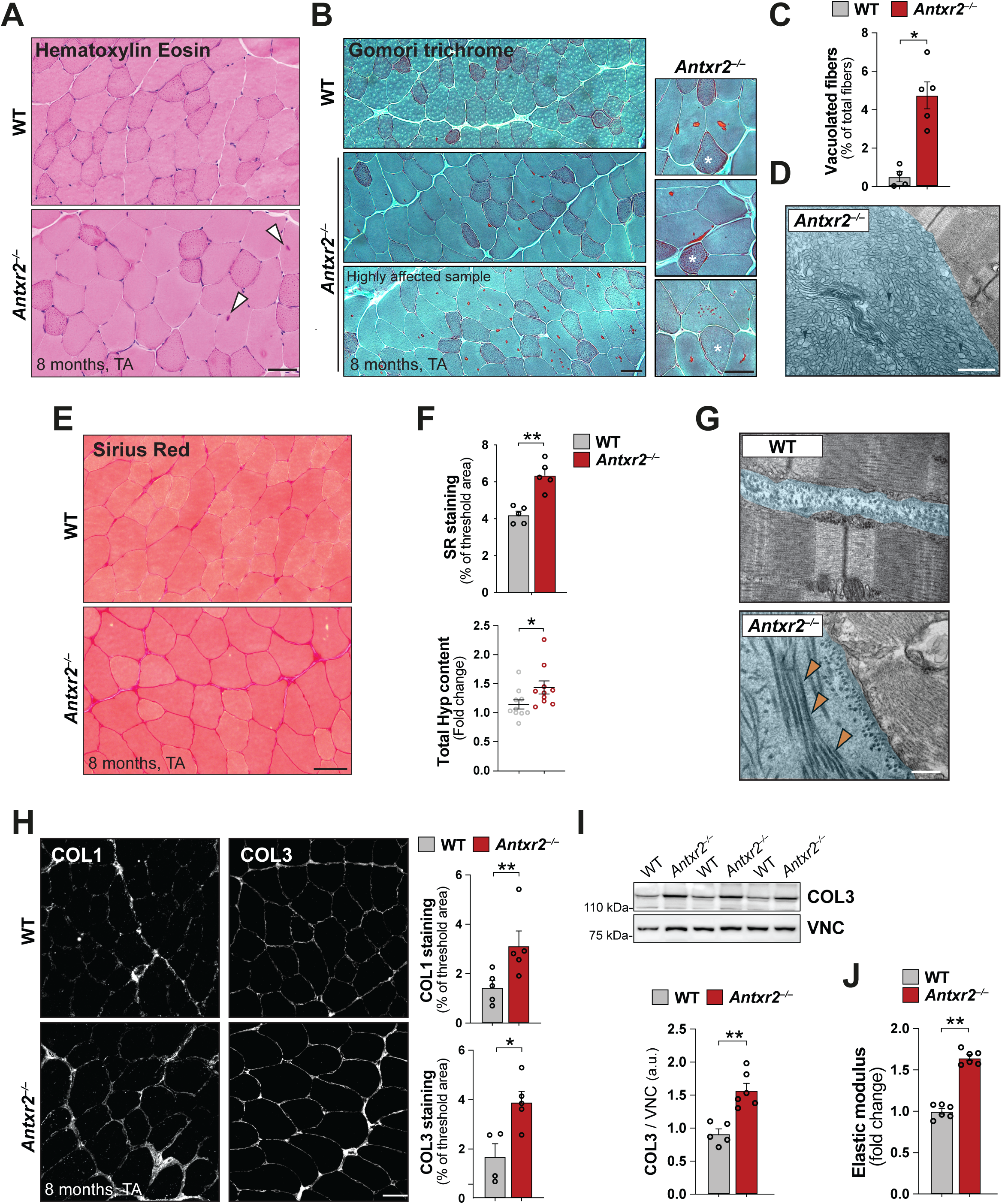
Muscles from adult ANTXR2-deficient mice display changes in myofibers and in the ECM. (**A**) Representative images of cross-sections of fresh frozen TA muscles from 8-month-old WT and *Antxr2*^−/−^ mice following staining with hematoxylin-eosin. White arrowheads indicates cytoplasmic inclusions. Scale bar, 50 μm. (**B**) Modified Gomori trichrome staining of cross-sections of fresh frozen TA muscles from 8-month-old WT and *Antxr2*^-/-^ male mice. Gomori-positive cytoplasmic inclusions (red), corresponding to tubular aggregates, are present in *Antxr2*^-/-^ muscles but not in WT muscles. The panels on the right show higher magnification images of *Antxr2*^-/-^ muscles, displaying different morphologies of tubular aggregates (intermyofibrillar, subsarcolemmal and fragmented). White asterisks indicate smaller oxidative fibers. Scale bar, 50 μm. (**C**) Quantification of the percentage of vacuolated myofibers, determined by modified Gomori trichrome staining as in (B). Data are expressed as mean ± s.e.m. (*n* = 4-5, each group; *, *P* <0.05; two-tailed Mann-Whitney test). (**D**) Representative transmission electron microscopy images of ultrathin sections of TA muscles from 8-month-old *Antxr2*^−/−^ male mice. Areas displaying tubular aggregates are highlighted in light blue. Scale bar, 1 μm. (**E**) Representative images of cross-sections of fresh frozen TA muscles from 8-month-old WT and *Antxr2*^−/−^ mice following staining with Sirius Red. Scale bar, 50 μm. (**F**) Top panel, histograms showing the percentage of threshold area positive for Sirius Red (SR, acquired under polarized light). Data are expressed as mean ± s.e.m. (*n* = 5, each group; **, *P* <0.01; two-tailed Mann-Whitney test). Bottom panel, Tukey box-and-whisker plots showing the total hydroxyproline (Hyp) content of gastrocnemius muscles from 8-month-old WT and *Antxr2*^−/−^ mice (*n* = 10, each group; *, *P* <0.05; two-tailed Mann-Whitney test). (**G**) Representative transmission electron microscopy images of TA ultrathin sections from 8-month-old WT and *Antxr2*^−/−^ mice. The endomysial area is highlighted in light blue. An abnormal accumulation of fibrillar collagens (orange arrowheads) is clearly visible in *Antxr2*^−/−^ muscles. Scale bar, 500 nm. (**H**) Representative confocal images of COL1 (white, left panels) and COL3 (white, right panels) immunostaining in cross-sections of TA muscles from 8-month-old WT and *Antxr2*^−/−^ mice. The histograms on the right show the percentage of threshold area positive for COL1 (top panel) and COL3 (bottom panel) staining. Data are expressed as mean ± s.e.m. (*n* = 4–5, each group; *, *P* <0.05; **, *P* <0.01; two-tailed Mann-Whitney test). Scale bar, 50 μm. (**I**) Western blotting (upper panel) and relative densitometric quantification (lower panel) of COL3 content in protein extracts from TA muscles of 8-month-old WT and *Antxr2*^−/−^ mice. Vinculin (VNC) was used as a loading control. Data are expressed as mean ± s.e.m. (*n* = 4–5, each group; **, *P* <0.01; two-tailed Mann-Whitney test). (**J**) Histogram showing the quantification of the elastic modulus of fresh TA muscles from 8-month-old WT and *Antxr2*^−/−^ mice. Data are expressed as mean ± s.e.m. (*n* = 6, each group; **, *P* <0.01; two-tailed Mann-Whitney test).

Since biochemical and immunofluorescent analysis of *Antxr2^-/-^*skeletal muscle revealed a progressive buildup of COL6, we next investigated whether this was accompanied by alterations in other ECM components. Sirius red staining showed increased deposition of fibrillar collagens in the endomysium of TA muscles from 8-month-old *Antxr2^-/-^* mice when compared to age-matched WT mice (Fig. 3E, F), a difference that was not observed in younger mice (Supplementary Fig. 4A, B). The increase in fibrillar collagens at 8 months, but not earlier, in TA muscles from *Antxr2^-/-^* mice was confirmed biochemically using hydroxyproline quantification, an amino acid signature for these ECM structures (Fig. 3F and Supplementary Fig. 4B).

Finally, the increased abundance of collagen fibrils in TA muscles was readily detectable by electron microscopy analysis of longitudinal ultrathin sections, in the vicinity of the myofiber basal lamina of ANTXR2-deficient muscles (Fig. 3G).

We next specifically monitored Collagen I (COL1) and Collagen III (COL3), the two most abundant fibrillar collagens in skeletal muscle, using immunofluorescence and western blotting. Both COL1 and COL3 were increased in TA and Gastrocnemius muscles of 8-month-old *Antxr2^-/-^* animals when compared to age-matched WT samples (Fig. 3H, I and Supplementary Fig. 4C). An increased abundance of these fibrillar collagens in TA muscles of *Antxr2^-/-^* mice was already apparent at 3 months of age (Supplementary Fig. 4D). These data indicate that the progressive extracellular accumulation of collagens in muscles lacking ANTXR2 extends to COL1 and COL3, albeit it seems to occur subsequently to COL6 accumulation during postnatal growth and maturity. Interestingly, the ECM changes elicited by ANTRX2 ablation do not appear to impinge on muscle basal lamina, i.e. the specialized innermost ECM of muscle endomysium in contact with the plasma membrane of myofibers. Indeed, immunofluorescence for Perlecan (HSPG2), Collagen IV (COL4) and Fibronectin (FN1), three major components of muscle basal lamina^35^, appeared similar in their amount and localization in muscles of 3-and 8-month-old WT and *Antxr2^-/-^*mice (Supplementary Fig. 5A-C).

To determine whether the accumulation of fibrillar collagens was associated with an increased expression of their genes, we quantified by RT-qPCR the mRNA levels for *Col1a1* and *Col3a1*. No significant differences in COL1 and COL3 expression were observed between 8-month-old WT and *Antxr2^-/-^* TA (Supplementary Fig. 4E), nor between sorted FAPs isolated from 8-month-old WT and *Antxr2^-/-^* muscles (Supplementary Fig. 4F). Moreover, TA and FAPs did no display any significant difference in the expression of the pro-fibrogenic *Acta2* and *Lox* genes between WT *Antxr2^-/-^* animals (Supplementary Fig. 4E, F). Therefore, the accumulation of ECM observed follows a mechanism that differs from classical fibrosis, where increases in the expression of collagen genes are observed^32^.

A typical consequence of increased collagen deposition is tissue stiffening^36,37^. We therefore measured the micromechanical properties of fresh skeletal muscle through a uniaxial tensile test, and found that 3-and 8-month-old *Antxr2^-/-^*TA muscles displayed a significantly higher elastic modulus when compared to the corresponding WT muscles, a clear sign of increased stiffness (Fig. 3J and Supplementary Fig. 4G).

Taken together, these data indicate that ANTXR2 deficiency triggers a progressive buildup of COL6 and fibrillar collagens in the interstitial ECM and increased muscle stiffness, without any evidence of increased expression of their genes, highlighting a different mechanism of ECM alteration than the typical fibrotic process.

### ANTXR2-deficient muscles develop progressive hypertrophy

We next assessed whether the ECM alterations detected in ANTXR2-deficient muscles affect myofiber organization. *Antxr2^-/-^* mice progressively developed heavier muscles than WT mice, a difference that became evident at 8 months of age, despite similar whole body weight (Fig. 4A and Supplementary Fig. 6A). Morphometric analyses of TA cross-sections showed a lower myofiber density (Fig. 4B, C), a significantly increased myofiber mean cross-sectional area (CSA; Fig. 4D, left panel) and mean minimum Feret diameter (MFD; Supplementary Fig. 6B, left panel) than age-matched WT. The difference became even more apparent when comparing the distribution of myofiber size (Fig. 4D and Supplementary Fig. 6B, right panels). Differences in CSA and MFD distribution were already apparent in 3-month-old mice, despite similar mean values (Supplementary Fig. 6C, D).

**Figure 4.**
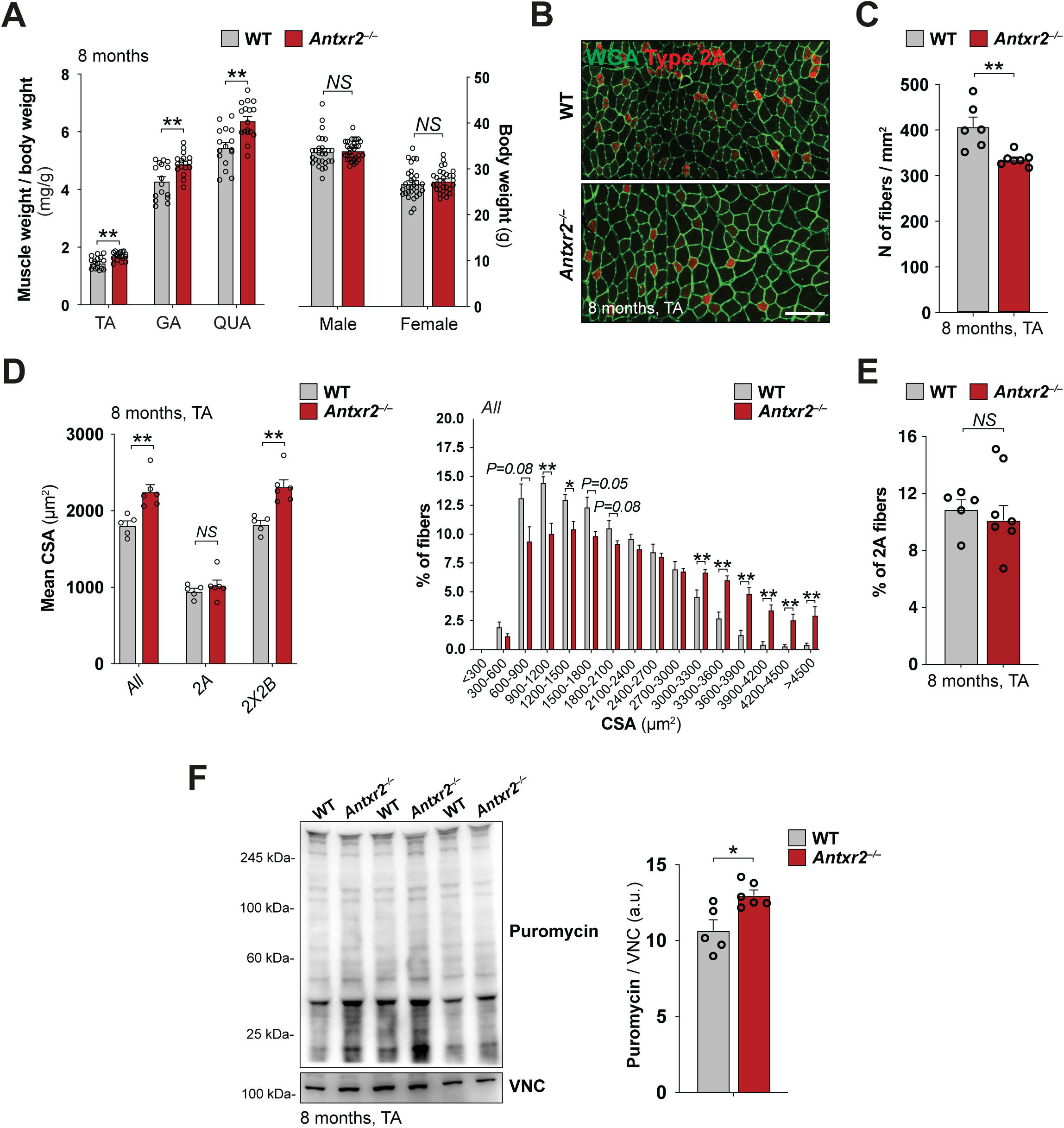
Muscles from adult ANTXR2-deficient mice are hypertrophic. (**A**) Normalized weight of TA, gastrocnemius (GA) and quadriceps (QUA) muscles (left panel) and total body weight (right panel) of 8-month-old WT and *Antxr2*^−/−^ mice. The weight of right and left muscles was averaged and normalized to body weight. Data are expressed as mean ± s.e.m. (*n* = 16–31, each group; NS, not significant, **, *P* <0.01; two-tailed Mann-Whitney test). (**B**) Representative confocal images of wheat germ agglutinin (WGA, green) and type 2A myofibers (positive for Myosin IIA, red) in cross-sections of TA muscles from 3-month-old wild-type and *Antxr2*^−/−^ mice. Scale bar, 50 μm. (**C**) Histograms showing the myofiber density (number of myofibers per area unit) in TA muscles of 8-month-old WT and *Antxr2*^−/−^ mice. Data are expressed as mean±s.e.m. (*n* = 5–6, each group; **, *P* <0.01; two-tailed Mann-Whitney test). (D) Left panel, mean CSA of myofibers of all types (All), of myofibers positive for myosin IIA (2A) and of myofibers negative for myosin IIA (2X2B) in TA muscles of 8-month-old WT and *Antxr2*^−/−^ male mice. Right panel, distribution of myofibers with different CSA (displayed as the percentage of total myofibers) in TA muscles of 8-month-old WT and *Antxr2*^−/−^ mice. Data are expressed as mean ± s.e.m. (*n* = 5–6, each group; NS, not significant; *, *P* <0.05; **, *P* <0.01; two-tailed Mann-Whitney test). (**E**) Histogram showing the percentage of type 2A myofibers in TA muscles of 8-month-old WT and *Antxr2^−/−^* mice. Data are expressed as mean ± s.e.m. (*n* = 5-7, each group; NS, not significant; two-tailed Mann-Whitney test). (**F**) Western blotting (left panel) and relative densitometric quantification (right panel) showing puromycine incorporation by SUnSET assay in TA muscles of 8-month-old WT and *Antxr2*^−/−^ mice. Vinculin was used as a loading control. Data are expressed as mean ± s.e.m. (*n* = 5, each group; *, *P* <0.05; two-tailed Mann-Whitney test). a.u., arbitrary units; VNC, vinculin.

The above data indicated that the absence of ANTXR2 leads to myofiber hypertrophy. Such hypertrophy often involves a switch from small oxidative myofibers (type 2A fibers, positive for myosin IIA) to large glycolytic myofibers (type 2X and 2B fibers, negative for myosin IIA)^38^. Immunostaining of TA muscles from 8-month-old mice for Myosin IIA, however, showed that the ratio of 2A to 2X/2B myofibers was not altered in *Antxr2^-/-^* animals (Fig. 4E). The 2X/2B fibers in *Antxr2^-/-^* TA muscles had significantly increased mean CSA and MFD (Fig. 4D and Supplementary Fig. 6B). Thus, the progressively hypertrophic phenotype was triggered in the more glycolytic myofibers.

Since myofiber hypertrophy is generally associated with an increase in protein synthesis, we measured protein synthesis *in vivo* using the puromycin-based SUnSET assay^39^. Animals at 8 months of age were injected with puromycin and, following muscle tissue isolation, puromycin-labelled truncated peptides were detected by western blotting. Puromycin incorporation into nascent polypeptides was significantly higher in TA muscles of 8-month-old *Antxr2^-/-^* mice when compared to age-matched WT samples (Fig. 4F).

Altogether, these data show that skeletal muscles of ANTXR2-deficient mice undergo a progressive hypertrophy of glycolytic myofibers, associated with an increase in protein synthesis.

### COL6 buildup in the ECM is the driver of muscle alterations in *Antxr2^-/-^* mice

In past studies, we showed that female ANTXR2-deficient mice suffer from severe accumulation of ECM in the uterus and found that this phenotype could be fully reverted by the additional knockout of the *Col6a1* gene, restoring female fertility^11^. We tested whether ablation of COL6 expression would similarly revert the myofiber hypertrophy displayed by ANTXR2-deficient muscles. Quantifications of CSA and MFD in cross-sections of TA (Fig. 5A, B) and Quadriceps (Fig. 5C) muscles from 8-month-old *Antxr2^-/-^*::*Col6a1^-/^*^-^ mice showed that the mean CSA and MFD were significantly lower than for WT mice, in fact very similar to what can be observed for *Col6a1^-/-^* mice (Fig. 5A-C). This was apparent when analysing the distribution of fiber CSAs, particularly for the TA, with a clear shift to smaller cross-sectional areas for fibers from mice lacking COL6, irrespective of the presence of ANTXR2 (Fig. 5B, C). Thus, the progressive buildup of COL6 is pivotal for the myofiber hypertrophy displayed by *Antxr2^-/-^* muscles.

**Figure 5.**
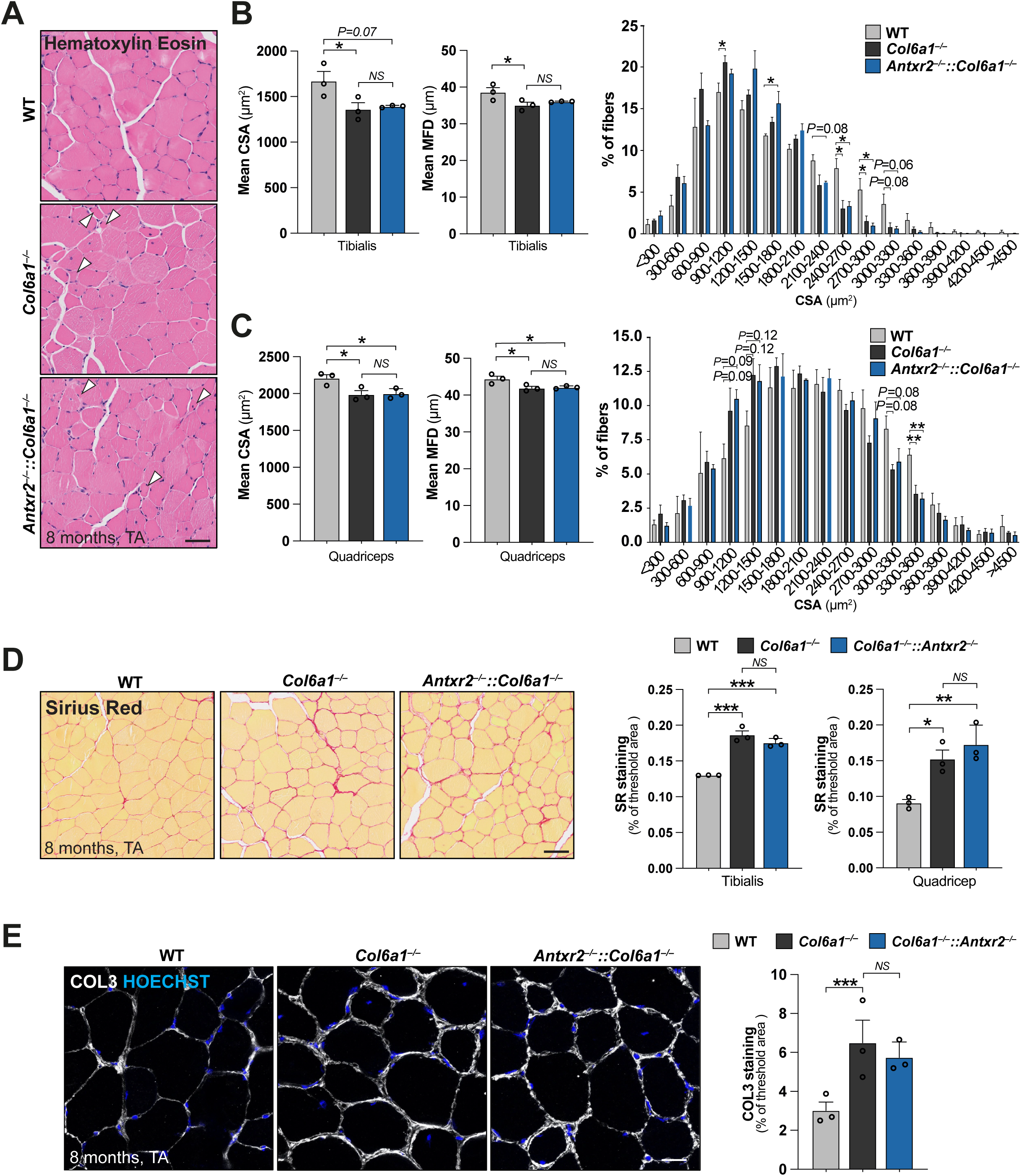
COL6 drives fibrillar collagen accumulation and myofiber hypertrophy in ANTXR2-deficient muscles. (**A**) Representative images of paraffin-embedded TA muscle cross-sections from 8-month-old WT, *Col6a1^-/-^* and *Antxr2^-/-^*::*Col6a1^-/-^*mice stained with hematoxylin-eosin (H&E). White arrowheads indicate small atrophic fibers. Scale bar, 50 μm. (**B**) Left panels, mean CSA and MFD of TA myofibers of 8-month-old WT, *Col6a1^-/-^* and *Antxr2^-/-^*::*Col6a1^-/-^* female mice. Right panel, distribution of myofibers with different CSA (displayed as the percentage of total myofibers) in TA muscles of 8-month-old WT, *Col6a1^-/-^* and *Antxr2^-/-^*::*Col6a1^-/-^*female mice. Data are expressed as mean ± s.e.m. (*n* = 3, each group; NS, not significant; *, *P* <0.05; one way ANOVA, with Dunnet’s test for multiple comparison). (**C**) Left panels, mean CSA and MFD of quadriceps myofibers of 8-month-old WT, *Col6a1^-/-^*and *Antxr2^-/-^*::*Col6a1^-/-^* male mice. Right panel, distribution of myofibers with different CSA (displayed as the percentage of total myofibers) in quadriceps muscles of 8-month-old WT, *Col6a1^-/-^* and *Antxr2^-/-^*::*Col6a1^-/-^* male mice. Data are expressed as mean ± s.e.m. (*n* = 3, each group; NS, not significant; *, *P* <0.05; \*\**P* <0.01; one way ANOVA, with Dunnet’s test for multiple comparison). (**D**) Left panel, representative images of paraffin-embedded TA muscle cross-sections from 8-month-old WT, *Col6a1^-/-^* and *Antxr2^-/-^*::*Col6a1^-/-^* mice stained with Sirius Red (SR). Right panel, Histograms showing the percentage of threshold area positive for Sirius Red in cross-sections of TA (left histogram) and quadriceps (right histogram) muscles from 8-month-old WT, *Col6a1^-/-^* and *Antxr2^-/-^*::*Col6a1^-/-^*mice. Data are expressed as mean ± s.e.m. (*n* = 3, each group; NS, not significant; *, *P* <0.05; **, *P* <0.01; ***, *P* <0.001; one way ANOVA, with Dunnet’s test for multiple comparison). Scale bar, 50 μm. (**E**) Representative confocal images of COL3 (white, left panels) immunostaining in TA cross-sections from 8-month-old WT, *Col6a1^-/-^* and *Antxr2^-/-^*::*Col6a1^-/-^* mice. The histograms on the right show the percentage of threshold area positive for COL3 staining. Data are expressed as mean ± s.e.m. (*n* = 3, each group; NS, not significant; ***, *P* <0.001; one-way ANOVA, with Dunnet’s test for multiple comparison). Scale bar, 50 μm.

We also tested whether the excessive deposition of fibrillar collagens observed in *Antxr2^-/-^* muscles would be reverted by the absence of COL6. Sirius red staining of TA and Quadriceps (Fig. 5D) and COL3 immunostaining of TA (Fig. 5E) of 8-month-old *Antxr2^-/-^*::*Col6a1^-/^*^-^ mice showed a significantly increased deposition of fibrillar collagens in the endomysial ECM when compared to the corresponding WT samples. A similar increase was however observed in *Col6a1^-/-^*mice (Fig. 5D, E). Thus, both in the absence and upon abnormal buildup of COL6, fibrillar collagens increase in muscle ECM, with no enhanced effect induced by ANTXR2 ablation.

Altogether, these data indicate that the hypertrophic phenotype and the fibrillar collagen accumulation displayed by *Antxr2*^-/-^ muscles are not merely caused by the absence of ANTXR2 itself, nor the increase in fibrillar collagens, but by the progressive buildup of COL6.

### ANTXR2-deficient mice have impaired motor function

Finally, we submitted mice to a battery of behavioural and physical tests. Importantly, *Antxr2^-/-^* and age-matched WT mice had similar voluntary activity during both nocturnal and diurnal, at 3-and 8-months (Fig. 6A and Supplementary Fig. 7A). To specifically evaluate muscle function, we determined plantar flexor muscle force by measuring torque production after electrical stimulation of the sciatic nerve in anaesthetized 8-month-old mice mice^40^. Of note, as for the muscles analysed above, the plantar flexor muscles of ANTXR2-deficient mice showed hypertrophy (Supplementary Fig. 7B). Surprisingly, at tetanus, the mice, whether *Antxr2^-/-^* or WT, showed similar absolute or normalized force (Fig. 6B and Supplementary Fig. 7C, D). Thus, the displayed muscle hypertrophy did not translate into an increase in strength. Interestingly, 8-month-old *Antxr2^-/-^* mice showed enhanced maximal force transmission speed, particularly at lower stimulation frequencies (Supplementary Fig. 7E) and a trend toward slower relaxation time after a single twitch (Supplementary Fig. 7F, G). These altered parameters may result from the increased stiffness of ANTXR2-deficient muscles, which could reduce the in-series compliance of a contracting muscle.

**Figure 6.**
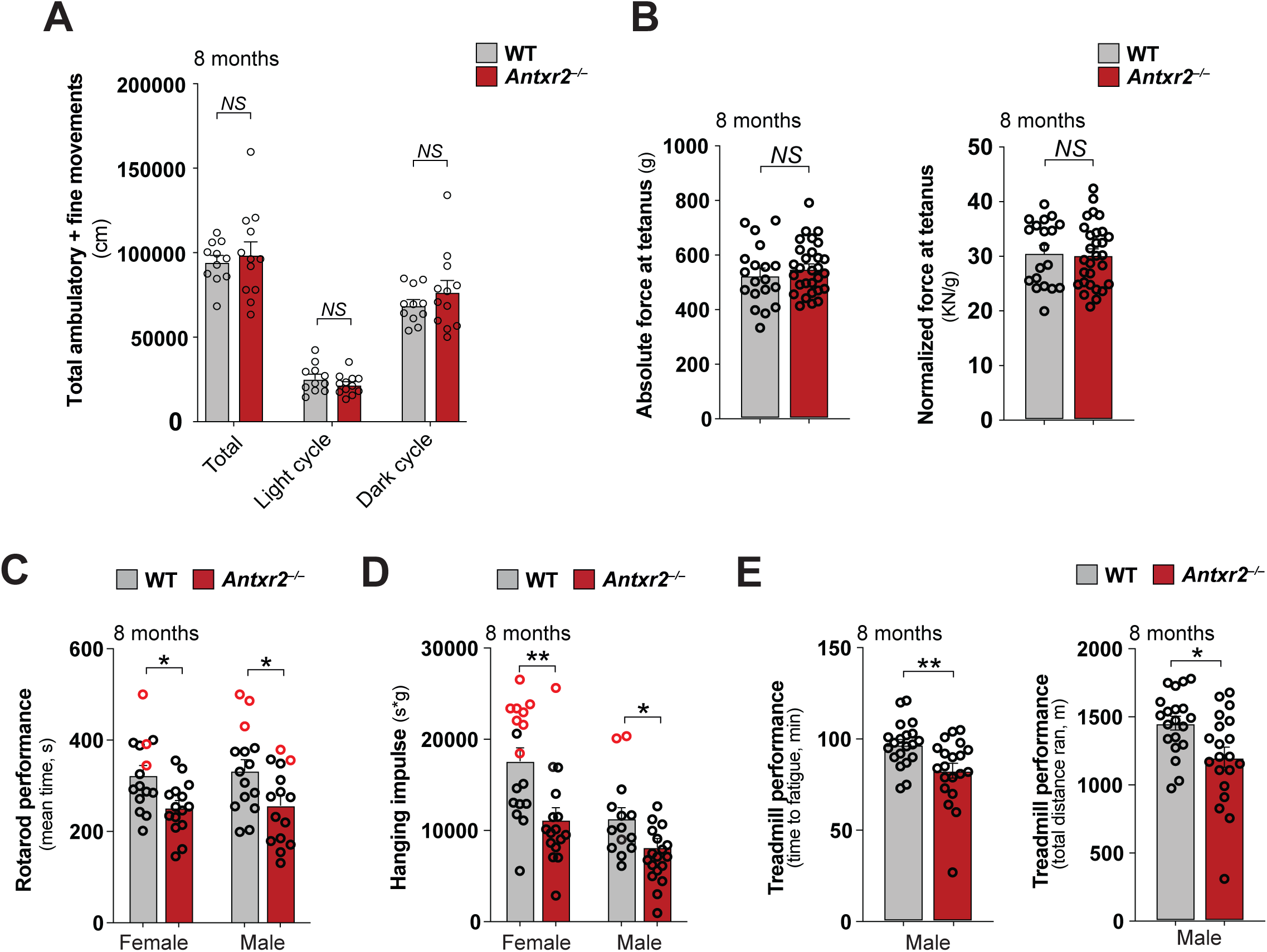
***Antxr2*^-/-^ mice display multiple signs of muscle weakness**. (**A**) Quantification of locomotor activity of 8-month-old male WT and *Antxr2*^-/-^ mice, as determined by 24-h-long home cage monitoring. The histograms show the sum of total ambulatory and fine movements during the entire day (Total) or only during the diurnal (Light cycle) or nocturnal (Dark cycle) periods. Data are expressed as mean ± s.e.m. (*n* = 11-12, each group; NS, not significant; two-tailed Mann-Whitney test). (C) Histograms showing the absolute and normalized force in plantar flexor muscles of 8-month-old WT and *Antxr2*^-/-^ male mice (*n* = 19-29 muscles from 10 WT and 15 *Antxr2*^-/-^ mice, respectively; NS, not significant; *, *P* <0.05; ** *P* <0.01; ***, *P* <0.001; two-tailed Mann-Whitney test). (**C**) Quantification of the rotarod performance test of 8-month-old WT and *Antxr2*^-/-^ female and male mice. Histograms represent the mean latency time. Red dots indicate the animals that reached the maximum time (500 s) in at least one of the two measurements. Data are expressed as mean ± s.e.m. (*n* = 15, each group; *, *P* <0.05; two-tailed Mann-Whitney test). (**D**) Quantification of the four-limb hanging performance of 8-month-old WT and *Antxr2*^-/-^ female and male mice. Histograms represent the holding impulse (i.e. best holding time normalized for body weight of the animal). Red dots indicate the animals that reached the maximum holding time (600 s) in at least one of the two evaluation tests. Data are expressed as mean ± s.e.m. (*n* = 14–17, each group; *, *P* <0.05; \*\**P* < 0.01; two-tailed Mann-Whitney test). (**E**) Quantification of the treadmill exercise performance of 8-month-old WT and *Antxr2*^-/-^ male mice. Histograms represent the time to fatigue (left panel) and the total distance run (right panel). Data are expressed as mean ± s.e.m. (*n* = 20, each group; *, *P* <0.05; **, *P* <0.01; two-tailed Mann-Whitney test).

We next investigated the motor coordination and tolerance to fatigue of 8-month-old male and female mice. *Antxr2^-/-^* mice showed a significantly lower performance when subjected to rotarod (Fig. 6C) and four-limb hanging (Fig. 6D) tests, differences that were not seen in younger mice (Supplementary Fig. 7H, I). Then, mice were further subjected to a treadmill running protocol. At the age of 8-months, when forced to run until they reached exhaustion, *Antxr2^-/-^*animals performed worse then age-matched WT animals, showing a decline in both the time needed to reach maximum exercise capacity and the total distance covered (Fig. 6E). This difference was not observed in younger mice, further emphasizing the progressive nature of the alterations induced by ANTRX2 deficiency (Supplementary Fig. 7J).

Altogether, these results indicate that ANTXR2-deficient mice undergo a progressive impairment of locomotor activity and a decreased physical fitness during adult life.

## DISCUSSION

Mutations in *COL6* genes that impair COL6 deposition lead to myopathy, highlighting the essential role of this non-fibrillar collagen in maintaining the proper architecture and function of the extracellular matrix^24^. While the pathological consequences of COL6 deficiency in skeletal muscle have been well documented, less is known about the importance of tuning the proper abundance of COL6 in the ECM. The amount of a given protein in the ECM is the result of biogenesis and turnover, with the first depending largely on gene expression and the second on degradation. ECM protein turnover is thought to occur primarily through extracellular processing by matrix metalloproteases (MMPs), a class of secreted, broad-substrate, Zn^2+^-dependent endopeptidases^5^. An alternative route for ECM and collagen remodelling is however emerging, where degradation occurs intracellularly following receptor-mediated endocytosis and transport to lysosomes^9,10^. This route is particularly relevant to COL6 given its relative resistance to MMPs^41–44^, possibly due to its distinctive structure, characterized by a somewhat short triple-helical region and large globular domains. While several receptors have been found to bind COL6^17^, ANTXR2 appears unique in its ability to mediate cellular uptake of COL6 and targeting to lysosomes^11^.

In this study, we therefore examined the role of ANTXR2 in skeletal muscle and found that it is predominantly expressed by interstitial FAPs, with negligible expression by mature myofibers. *In vivo* ANTXR2 ablation in *Antxr2^-/-^* mice results in the early and progressive COL6 buildup in intramuscular ECM, leading to elevated amounts of fibrillar collagens and increased stiffness in adult mice. These ECM changes are accompanied by myofiber changes, with a non-functional hypertrophy and clearly identifiable signs of functional impairment and muscle weakness. These findings underline that proper control of extracellular COL6 levels — through both gene expression and ANTXR2-mediated degradation — is crucial for maintaining muscle function, as both deficiency and excess of COL6 are detrimental.

*Antxr2^-/-^* mice appear normal at birth and shortly after. At four weeks, they start a progressive accretion of extracellular COL6 in the intramuscular connective tissue. This indicates that, under physiological conditions, ANTXR2-mediated COL6 degradation needs to occur already in young mice to ensure proper homeostasis of this key muscle ECM component. The progressive COL6 buildup in muscles of *Antxr2^-/-^* mice is paralleled by an ensuing accumulation of fibrillar collagens, which is particularly prominent at 8 months of age. In contrast, the basal lamina of ANTXR2-deficient muscles shows no apparent sign of alterations, indicating that defects are restrained to the outer part of the intramuscular connective tissue layers.

In healthy muscle, the ECM is most synthesized and remodelled by FAPs^36^. Interestingly, both COL6 and ANTXR2 are abundantly expressed by FAPs, indicating that these cells both produce and degrade COL6. Of note, ablation of *Antxr2* does not lead to upregulation of fibrotic markers, such as *Acta2* and *Lox*, neither in muscles nor in FACS-sorted FAPs. Therefore, the progressive accumulation of COL6 does not trigger myofibroblast differentiation. Consistently, we did not detect other typical signs of fibrotic changes, such as fibrous-adipose replacement and cellular infiltration^32^. This is further supported by electron microscopy imaging, which, while revealing a pronounced accumulation of fibrillar collagens in the intramuscular connective tissue of adult *Antxr2^-/-^* mice, does not show the chaotic and irregular arrangement of collagen fibrils typical of a fibrotic condition. Taken together, these data indicate that the lack of either COL6 or ANTXR2 leads to increased fibrillar collagens in the ECM, but the processes underlying such ECM changes involve distinct mechanisms. Both mice and patients with COL6 deficiency display upregulation of genes coding for COL1 and COL3, consistently with the presence of muscle fibrosis^45,46^. In the absence of ANTXR2, however, expression of collagen genes is unchanged, but COL6 accumulates due to altered degradation, and as a secondary effect fibrillar collagens accumulate in the ECM, most likely due to the ability of COL6 microfilaments to tightly interconnect with collagen fibrils^47–51^.

Our data on the biomechanical properties of TA muscles from adult *Antxr2^-/-^* mice indicate that the progressive accumulation of COL6 and fibrillar collagens in the ECM is paralleled by significant changes in the elastic modulus, with increased muscle stiffness. Interestingly, this also occurs in muscles of 3-month-old *Antxr2^-/-^* animals, when COL6 – but not fibrillar collagens – already display increased ECM deposition. This observation supports earlier findings indicating that COL6 contributes to the fine regulation of tissue stiffness, affecting the mechanical properties of the tissue^47,48,52,53^.

Although ANTXR2 expression by mature myofibers is very low, its ablation and the ensuing progressive accretion of COL6 in the ECM lead to a number of distinctive myofiber changes. First, myofibers of adult *Antxr2^-/-^*male mice consistently show the presence of intracellular membranous structures positive for Gomori staining and with features of tubular aggregates. Such tubular aggregates represent myofiber inclusions consisting of tubules derived from the sarcoplasmic reticulum, typically more abundant in males, found in a variety of muscle disorders characterized by weakness, myalgia and cramps, being also described in muscles undergoing age-related sarcopenia^33,34^.

The most remarkable change elicited by ANTXR2 ablation is however myofiber hypertrophy, associated with increased protein synthesis. This hypertrophic phenotype is linked to the progressive COL6 buildup, since muscles of *Antxr2^-/-^*::*Col6a1^-/^*^-^ double knockout mice showed signs of myofiber atrophy, rather than hypertrophy, akin to what is displayed by muscles of *Col6a1^-/-^* mice^40^. Notably, the myofiber hypertrophy displayed by ANTXR2-deficient muscles does not yield increased force, leading instead to functional defects, as revealed by various behavioural tests and *in vivo* quantification of muscle contractile force showing that *Antxr2^-/-^* mice experience impaired locomotor activity and reduced physical fitness when physically challenged. Taking into account that myofibers of adult WT muscles barely express ANTXR2, these results strongly supports the concept that alterations in ANTXR2-deficient FAPs, in particular their inability to internalize COL6, result in ECM buildup in *Antxr2^-/-^* muscles, in turn leading to increased muscle stiffness and compromised lateral transmission of force between myofibers and from muscle to tendons, especially during high-intensity exercise.

In conclusion, our data indicate that ANTXR2 is required for the structural integrity and function of skeletal muscle. In particular, it contributes to the regulation of the extracellular abundance of COL6 and thereby to the proper organization and dynamics of the ECM. As such, ANTRX2 represents an ECM regulator that may have significant implications in various physiological and pathological contexts where ECM remodelling occurs, such as aging and fibrosis. Loss-of-function mutations of the human *ANTXR2* gene lead to a devastating rare genetic disease, Hyaline Fibromatosis Syndrome (HFS)^11,15^. Consistent with the here identified role of ANTXR2 in muscle functions, HFS patients commonly experience muscle stiffness, weakness and joint contractures^54^. In particular, prominent skeletal muscle involvement with myopathic changes was reported in some infantile patients^55,56^. Thus, our findings indicate that *Antxr2^-/-^* mice also represent a valuable model for investigating the pathomolecular mechanisms underlying muscle alterations in HFS.

## MATERIAL AND METHODS

### Mouse maintenance

Age-matched WT, ANTXR2 knockout (*Antxr2*^-/-^), COL6 knockout (*Col6a1*^-/-^)^57^ and ANTXR2/COL6 double knockout (*Antxr2^-/-^*::*Col6a1^-/-^*) mice were maintained in the C57BL/6J background and housed in specific pathogen-free controlled environment (23° C temperature, 12 h light/12 h dark cycle). Mice were provided with *ad libitum* standard chow and water. *Antxr2^-/-^* mice were generated by targeted deletion of exon 3 of the *Antxr2* gene^11^, while *Antxr2^-/-^*::*Col6a1^-/-^*double knockout mice were generated by crossing *Antxr2*^+/–^ and *Col6a1*^+/–^ heterozygous animals. According to the experiment, 5-day-old pups or 4-, 12-and 35-week-old male and female mice were used. All procedures were performed according to protocols approved by the Veterinary Authorities of the Canton Vaud (licences VD 2144.4 and VD 3918) and by the Ethics Committee of the University of Padova (OPBA, n. 100/2020-PR) and in agreement with the Swiss and Italian pertinent laws.

### Hindlimb muscle dissection

Hindlimb skin and connective tissue (epimysium) were gently removed to expose the underlying tibialis anterior (TA), quadriceps femoris and gastrocnemius muscles. For 5-day-old pups, given the small size of skeletal muscles and the technical challenges of a proper tissue dissection, only the gastrocnemius muscle was collected and analyzed. Once harvested, muscles were quickly immersed in an isopentane-containing beaker, precooled by apposition to liquid nitrogen vapours. Frozen muscles were then stored at-80° C until subsequent analyses. Where indicated, extensor digitorum longus (EDL) muscles were collected and immediately fixed with 4% paraformaldehyde for 60 min at 4° C, washed twice in phosphate-buffered saline (PBS) and stored in PBS at 4° C. Before immunofluorescent analysis, EDL muscles underwent mechanical dissociation using scalpels to isolate either single myofibers or myofiber bundles.

### Histological analyses

Cross-sections (10-μm-thick) of frozen muscles were used for the following staining protocols. For hematoxylin-eosin staining, slides were hydrated in bidistilled water for 2 min and stained with haematoxylin (Sigma-Aldrich) for 1-2 min, followed by washing with bidistilled water for 3 min. The slides were incubated for 1 min with eosin (Sigma-Aldrich), and subsequently rinsed in bidistilled water. For sirius red staining, slides were incubated in 4% paraformaldehyde for 15 min at room temperature, washed twice in PBS and incubated with 300 μL of picrosirius red solution (Sigma-Aldrich) in a moist chamber for 1 h at room temperature. Slides were then washed four times for 5 min in 0.5% glacial acetic acid (Sigma-Aldrich) under gentle shaking. For modified Gömöri trichrome staining, slides were hydrated in bidistilled water for 2 min and then incubated with Gomori trichrome (CliniSciences) for 10 min, and differentiated with 0.2% acetic acid for 10 min. After dehydration, slides were immersed in xylene and mounted with Eukitt Quick-hardening mounting medium (Sigma-Aldrich). Bright-field images were captured by using a Leica DM-R microscope equipped with a digital camera or an Olympus VS200 slide scanner. Sirius red staining images were acquired with a digital camera-equipped Leica DM6 B microscope under polarized light. Gomori-positive cytoplasmic inclusions were manually counted on the entire stained muscle section. Quantification of sirius red staining was performed with Fiji^58^.

### Immunofluorescence imaging

Cross-sections (10-μm-thick) of frozen muscles were fixed and permeabilized in cold 1:1 methanol-acetone solution for 10 min at - 20° C, washed twice in PBS under gentle shaking and saturated with 10% goat serum (GS) in PBS at room temperature. Single myofibers or myofiber bundles were permeabilized with 0.2% Triton X-100 in 10% GS for 20 min and finally saturated with 10% GS in PBS at room temperature. After 60 min, primary antibodies were prepared in 5% GS and incubated overnight at 4° C. The following primary antibodies were used: rat monoclonal anti-ANTXR2 (1:50; clone 8F7, homemade produced^11^); rabbit recombinant anti-Vimentin (1:200; D21H3, Cell-signaling technology); rabbit polyclonal anti-COL6 α3 (1:100; kindly provided by Dr. Raimund Wagener, Cologne^59^); guinea pig polyclonal anti-COL6 α6 (1:200; kindly provided by Dr. Raimund Wagener, Cologne)^59^; rabbit polyclonal anti-COL1 α1 (1:200; BP2201, Acris); rabbit polyclonal anti-COL3 α1 (1:200; 22734-1-AP, Proteintech); rabbit anti-Laminin (1:800; L9393, Sigma-Aldrich); rabbit polyclonal anti-FN1 (1:200; F3648, Sigma-Aldrich); rat monoclonal anti-HSPG2 (1:200; sc-33707, Santa Cruz Biotechnology); rabbit polyclonal anti-COL4 (1:200; AB756P, Millipore); mouse monoclonal anti-Myosin Heavy Chain IIA (1:100; SC-71, Developmental Studies Hybridoma Bank); rabbit polyclonal anti-Calsequestrin (1:200; PA1-913, Thermo-Fisher); mouse monoclonal anti-Stromal Interaction Molecule 1 (STIM1, 1:100; 610954, BD Transduction Laboratories); rabbit polyclonal anti-Calcium Release-Activated Calcium Modulator 1 (CRACM1/Orai1, 1:100; 4281, ProSci inc.); mouse monoclonal anti-Dihydropyridine Receptor (DHPR, 1:100; clone MA3-920, Sigma-Aldrich); mouse monoclonal anti-Sarco/Endoplasmic Reticulum Calcium ATPases (SERCA1/2/3, 1:100; sc-271669, Santa Cruz Biotechnology). The next day, slides or fiber bundles were washed three times in PBS for 10 min under gentle shaking and then incubated with appropriate secondary antibodies (Jackson ImmunoResearch) and nuclear dye Hoechst 33258 for 1 h at room temperature. To stain F-Actin, Phalloidin conjugated with Alexa Fluor 647 (Thermo-Fisher) was added together with secondary antibodies. Slides and myofiber bundles were washed three times in PBS for 10 min under gentle shaking at room temperature, and mounted in ProLong Glass Antifade Mountant (Invitrogen). Images were acquired with a Leica SP8 Stellaris laser-scanning confocal microscope. Quantification of the thresholded area was performed with Fiji.

### RNA *in situ* hybridization

RNA *in situ* hybridizations were performed using the RNAscope Fluorescent Multiplex V2 Assay (Bio-Techne, cat. n. 323110) following the manufacturer’s instructions. Briefly, 10-μm-thick cross-sections of TA were fixed with 4% paraformaldehyde for 60 min at 4° C, washed twice with PBS, dehydrated and quenched with hydrogen peroxide for 10 min at room temperature. Then, slides were washed twice in PBS and treated for 10 min at room temperature with protease IV. Probes for mouse *Antxr2* (Bio-Techne, cat. n. 468651) were incubated for 2 h and RNAscope amplification was performed following the manufacturer’s instructions using Opal 650 fluorophore reagent pack (Akoya Biosciences, cat. n. FP1496001KT). Coupled immunofluorescence analysis was performed as described above without the permeabilization step. Finally, slides were mounted in Prolong Gold antifade mounting medium (Thermo-Fisher) and imaged using a Leica SP8 laser-scanning confocal microscope. Quantification of mRNA dots was performed with QuPath^60^.

### Muscle morphometric analysis

Cross-sections (10-μm-thick) of frozen TA muscles were incubated with 1 μg/mL of wheat germ agglutinin (WGA) conjugated with Alexa Fluor 488 (Invitrogen) and 2.5 μg/mL of Hoechst 33258 (Sigma-Aldrich) in PBS for 20 min at room temperature. Slides were washed twice in PBS for 5 min under gentle shaking and mounted in 80% glycerol in PBS. For each sample, the entire stained section was acquired with a Leica SP8 Stellaris laser-scanning confocal microscope (Leica Microsystems). Myofiber cross-sectional area (CSA) and minimum Feret diameter (MFD) were measured with the Myosoft plugin^61^.

### Transmission electron microscopy

TA muscles were fixed overnight at 4° C with 2.5% glutaraldehyde in 0.1 M cacodylate buffer, washed with 0.1 M cacodylate buffer, post-fixed for 2 h with 1% osmium tetroxide, and embedded in Epon812 (Electron Microscopy Sciences). Transversal and longitudinal ultrathin sections were stained with uranyl acetate and lead citrate and observed with a Philips EM400 electron microscope operating at 100 kV.

### Western blotting

Frozen muscles were pulverized by grinding in liquid nitrogen and lysed in extraction buffer (50 mM Tris-HCl, pH 7.5; 150 mM NaCl; 10 mM MgCl_2_; 1 mM EDTA; 10% glycerol; 0.5 mM DTT; 2% SDS; 1% Triton X-100) supplemented with protease and phosphatase inhibitors (Sigma-Aldrich). SDS-insoluble fractions were obtained after centrifuging the protein extracts and lysing the pellet in urea buffer (8 M urea, 2% SDS) supplemented with protease inhibitors (Sigma-Aldrich). Protein concentration was determined by the BCA Protein Assay Kit (Thermo-Fisher) and 30-40 μg of protein samples were used for SDS-PAGE in polyacrylamide Novex NuPAGE Bis-Tris 4-12% gels (Invitrogen). For native COL6 detection, protein samples were run in a homemade composite 2.5% acrylamide/0.5% agarose gel under non-reducing conditions for higher molecular weight separations^62^. Gels were electrotransferred onto polyvinylidene difluoride membranes (Millipore), stained with Ponceau S (Sigma-Aldrich) and blocked for 1 h with 5% non-fat dry milk in Tris-buffered saline solution containing 0.1% Tween 20 (TBS-T), followed by overnight incubation at 4° C with primary antibodies. The following primary antibodies were used: rabbity polyclonal anti-COL6 (1:1,000; 70-XR95, Fitzgerald Industries, for native COL6 detection); rabbit polyclonal anti-COL6 α1 (1:1,000; sc-20649, Santa Cruz Biotechnology); rabbit polyclonal anti-COL6 α3 (1:1,000; homemade produced, kindly provided by Dr. Raimund Wagener, Cologne)^59^; rabbit polyclonal anti-COL3 α1 (1:2,000; 22734-1-AP, Proteintech); mouse monoclonal anti-GAPDH (1:125,000; MAB374, Millipore); mouse monoclonal anti-Vinculin (1:1,000; V4505, Sigma-Aldrich); mouse monoclonal anti-puromycin (1:2,500; clone 12D10, Millipore). After three washes in TBS-T, membranes were incubated for 1 h with horseradish peroxidase-conjugated anti-rabbit or anti-mouse secondary antibodies (1:1,000; Bethyl Laboratories). Detection was carried out by SuperSignal West Pico (Thermo-Fisher) and densitometric quantification was performed by Fiji software.

### Hydroxyproline assay

For this assay the gastrocnemius muscle was selected based on its larger size, enabling the collection of sufficient tissue material required for the analyses. Muscles were pulverized, collected in a pre-cooled tube, weighted, and maintained over liquid nitrogen vapours. Next, 7 μL of HCl and 7 μL of bidistilled water were added for each mg of tissue, and the tubes were kept at 130° C for 3 h to dissolve the tissue sample. Lysates were further homogenized by pushing the specimens through a 26G needle four times and then centrifuged at 12,000 rcf for 10 min at room temperature. 30 μL of supernatant were transferred into a new tube and 625 μL of chloramine T buffer (Sigma-Aldrich) were added to each sample and incubated for 20 min at room temperature. Then, 625 μL of 4-(dimethylamino)benzaldehyde (DMAB) buffer (Sigma-Aldrich) was added and incubated for 10 min at 65° C. Reaction was stopped by putting tubes on ice and 200 μL of each sample were loaded in duplicate in a 96-well plate. Color intensity was measured with a microplate spectrophotometer reader (Tecan Life Sciences), by exciting the samples at a wavelength of 540 nm. A calibration curve obtained from known concentrations of hydroxyproline (Sigma-Aldrich) was used to extrapolate the concentrations of the samples from their experimental absorbances.

### *In vivo* protein synthesis

*In vivo* protein synthesis in skeletal muscle was measured using the SUrface SEnsing of Translation (SUnSET) technique^39^. Mice were weighed and subjected to injection with puromycin (Sigma-Aldrich) dissolved in 100 μL of sterile PBS at the concentration of 0.04 μmol/g body mass. 30 min after puromycin injection, TA muscles were collected, frozen in liquid nitrogen and stored (−80 °C) for western blot analysis.

### Fluorescence-activated cell sorting

FAPs were obtained from gastrocnemius muscle using a previously described protocol with slight modifications^63^. In brief, muscle tissue underwent enzymatic digestion with 2 mg/mL Collagenase II (Sigma-Aldrich), 1.5 U/mL Collagenase D (Sigma-Aldrich), 2.4 U/mL Dispase II (Sigma-Aldrich), and 250 mM CaCl_2_. Following digestion, muscles were gently mashed and supplemented with red blood cell lysis buffer (0.15 mM NH_4_Cl, 8.4 mM KHCO_3_, 1.2 mM EDTA in PBS) before filtration through 70 µM and 40 µM cell strainers to obtain single-cell suspensions. These suspensions were then incubated with primary antibodies targeting specific cell markers for 30 min at 4° C in a rotating wheel. The following antibodies were used: CD45-eFluor 450 (1:50; Invitrogen, clone 30-F11), CD31 (PECAM-1)-PE-Cy7 (1:50; Invitrogen, clone 390), CD140a (PDGFRa)-APC (1:25; Invitrogen, clone APA5), Ly6A/E (Sca-1)-FITC (1:50; Invitrogen, clone D7), Viability Fixable Dye 405/520 (1:100; Miltenyi Biotec). FAPs (CD45^-^/CD31^-^/PDGFRα^+^/Sca-1^+^) were subsequently isolated using flow sorting in a FACS Canto II (Becton Dickinson). Sorted FAPs were collected into cold PBS before RNA extraction.

### RT-qPCR

Muscle samples were pulverized by grinding in liquid nitrogen and total RNA was extracted using 0.5 mL TRIzol Reagent (Thermo-Fisher). RNA extraction from sorted FAPs was performed using the Nucleospin RNA XS isolation kit (Macherey-Nagel). RNA was quantified with Nanodrop ND-1000 (Nanodrop Technologies). Reverse transcription was performed with 500 ng total RNA and M-MLV Reverse Transcriptase (Thermo-Fisher), using random hexamers as previously described^64^. The resulting cDNAs were processed for quantitative real-time PCR using Rotor-Gene SYBR Green PCR Kit mastermix (Qiagen) and a Rotor-GeneQ thermocycler (Qiagen). Each sample was loaded in triplicate and analyzed with RotorGene Q 2.0.24 software. *Rps16* or *Gapdh* were used as housekeeping gene controls. Primers are listed in Supplementary Table 1.

### Quantification of muscle stiffness

The mechanical properties of fresh TA muscles were characterized through a tensile test using a uniaxial testing machine (ZwickRoell ProLine Z005) equipped with a 10 N load cell. Muscle samples were fixed to the clamps by using two suture needles with inextensible thread, suitably shaped to prevent the specimen from slipping. The test was performed by imposing a strain rate of 10% of the initial length (distance between the needles) per minute, and was stopped at sample failure, acquiring total displacement (mm) and standard force (N). Data were then analyzed in MS Excel^®^, obtaining strain (%) and stress (kPa) values and plotting the stress-strain curves. The elastic modulus was then determined by performing a linear regression on the linear portion of the curves.

### Spontaneous locomotor activity

Global tracking of the voluntary activity over 24 h, covering both nocturnal and diurnal periods was measured with a Comprehensive Laboratory Animal Monitoring System (CLAMS, Columbus Instruments). Mice were placed individually in the plexiglas chambers and the spontaneous activity, the food intake and the water intake were measured simultaneously. Ambulatory movements and fine movements were measured.

### Four-limb hanging test

Mice were placed on a grid, which was kept upside down above an empty cage filled with bedding materials. The hanging time, i.e., the time until the animal fell, was recorded. A fixed limit of 900 s was set, and each mouse was tested twice with a 30-min break between measurements. Holding impulse was obtained by multiplying the hanging time for the respective mouse body weight. The maximum holding time was used for the analysis according to TREAT-NMD standard operating procedures (DMD_M.2.1.005).

### Rotarod performance test

Two days before the experiments, mice underwent training sessions on a rotating rod (Ugo Basile) for two sessions lasting 10 min each, maintained at a constant speed of 7 rpm, with a 30-min break in between sessions. The day of the experiment, after 3 min warmup at a constant speed of 7 rpm, a ramp of 7-30 rpm in 300 s was initiated. The ending time point was set at 500 s, and mice that did not fall off within this period were given a maximum score. The test was repeated after 30 min and the average value was used for statistics.

### Exercise performance test

Before the test, mice were habituated to the treadmill (Panlab) for 10 min at a speed of 5 cm/s. Acute exercise was started at the very moderate speed of 15 cm/s with a 10° positive inclination, and the speed was gradually increased of 3 cm/s every 12 min. The run distance and the number of received shocks (0.1-0.3 mA) were monitored. If a mouse received more than five shocks in two consecutive minutes, it was considered as having reached exhaustion and placed back in its homecage.

### *In vivo* muscle force measurement

Mice were anesthetized with one i.p. injection of a solution containing 100 mg/kg ketamine and 10mg/kg xylazine and, a few minutes later, one i.p. injection of 5 mg/kg altadol as a pain relief treatment. A small incision was made from the knee to the hip and the sciatic nerve was exposed. Teflon-coated multistranded stainless-steel wires (AS 632; Cooner Wire Company, Chatsworth, CA, USA) was sutured with 5-0 silk thread at each side of the nerve and the wound was closed using the same thread. To avoid recruitment of the ankle dorsal flexors, and the consequential reduction of torque, the common peroneal nerve was cut. Torque production of the stimulated plantar flexors was measured using a muscle lever system (Model 305C; Aurora Scientific, Aurora, ON, Canada), data were acquired using the DMCv5.3 software (Aurora Scientific), and electrical stimulation was achieved using a Grass S11 stimulator (Grass Instrument Company, Quincy, MA, USA). The force-frequency curves were determined by stepwise increasing stimulation frequency (from single pulse to 20, 40, 55, 75, 100 and 150 Hz; train duration, 200 ms; single pulse width 500 µsec; 4-6 V), with a pause of 30 s between stimuli to avoid muscle fatigue. Time parameters (time to peak and half relaxation time) were measured on the force trace obtained with a single stimulus (twitch). Contraction parameters were obtained from the tetanus trace obtained at 100 Hz electrical stimulation. The fast rate of force development was measured as a linear fitting (max speed, g/ms). Normalized force was obtained by dividing absolute force to muscle weight (obtaining kN/g). Representative diagrams of twitch and absolute force traces are shown in **Fig. S7G**. Raw data analysis was performed using a custom-made automatic LabView software.

### Single-nucleus RNA sequencing dataset analysis

The GSE147127 dataset, containing single-nucleus RNA-seq data of skeletal muscles from C57BL/6 mice of different ages^28^, was used for this analysis. Normalized count matrix and dimensional reduction data were downloaded from https://www.synapse.org/#!Synapse:syn21676145/files/ as Seurat object. Seurat v4.3.0.1 was then used for plotting features of interest on the pre-computed UMAP^65^. The dotPlot function from scCustomize v1.1.3 was used to plot the percentage of cells expressing selected genes.

### Statistics

All results are provided as mean ± s.e.m. Statistical analysis of data was performed with an unpaired two-tailed Mann-Whitney test (GraphPad). For experiments with more than two conditions, one-way or two-way analysis of variants (ANOVA) was used (GraphPad). When ANOVA revealed significant differences, further analysis was carried out using Dunnet’s or Sidak’s multiple comparison test. Statistical significance was set at *P* <0.05. All histograms display individual values, and the number of biological replicates (always greater than three) as well as the specific statistical tests used are indicated in the figures’ captions.

## Acknowledgements

We thank Dr. Dario Bizzotto (University of Padova) and Dr. Audrey Chuat (EPFL) for the maintenance of mouse colonies and genotyping; Dr. Olha Novokhatska (EPFL) for the initial involvement in the locomotor activity tracking; Dr. Raimund Wagener (University of Cologne, Cologne, Germany) for providing COL6 antibodies; the core facilities BIOP and CPG at EPFL for sharing their equipment and expertise; Dr. Matilde Cescon (University of Padova) and all the members of G. van der Goot and P. Bonaldo labs for helpful discussions and suggestions. This work was funded by the Italian Ministry of University and Research (Grant 2022MXH3JY to P.B.), the Telethon Foundation (GGP19229 to P.B.), and the Swiss Foundation for Research on Muscle Diseases to F.G.v.d.G.. S.M. was supported by the European Molecular Biology Organization EMBO Scientific Exchange (Grant Number 9111) and by the Marie-Curie MSCA Global Fellowship (Grant agreement 101149789).

## Author contributions

S.M., F.G.v.d.G. and P.B. designed the study. S.M. performed most of the experiments and relative analyses. L.J.B., M.S., L.A., B.K., and L.G. contributed to *in vivo* experiments. J.D. analyzed transcriptomic data. L.N. and B.B. performed *in vivo* muscle force measurements. G.P. and N.F. contributed to hydroxyproline content analysis. P.S. performed transmission electron microscopy analysis. A.D.A., G.M.F., P.Br. and G.V. performed muscle stiffness measurements. S.M., F.G.v.d.G. and P.B. wrote the original draft and all the other authors contributed to the final version of the manuscript.

